# From Selfies to Science – Precise 3D Leaf Measurement with iPhone 13 and Its Implications for Plant Development and Transpiration

**DOI:** 10.1101/2023.12.30.573617

**Authors:** Gabriel Bar-Sella, Matan Gavish, Menachem Moshelion

## Abstract

Advanced smartphone technology now integrates sophisticated sensors, increasing access to high-precision data acquisition. This study tested the hypothesis that the iPhone 13-Pro camera, with LiDAR technology, can accurately estimate maize leaf surface area (Zea mays). 3D point cloud models enabled non-destructive data collection, and four methods for canopy area extraction were evaluated in relation to plant transpiration rates. Results showed a strong correlation (R^2^=0.92, RMSE=49.78) between manually scanned and iPhone-estimated plant surface areas. Additionally, the stem-to-plant surface area ratio was found to be 12.3% (R^2^=0.9, RMSE=28.42). Using this ratio to predict canopy area showed a significant correlation (R^2^=0.83) with actual canopy measurements. The iPhone’s surface area measurement tool offers an advantage by scanning the entire plant surface, unlike traditional leaf area index measurements, which often cannot penetrate the canopy. Moreover, real-size surface measurement of the canopy correlated strongly (R^2^=0.83) with whole canopy transpiration rates measured gravimetrically. This study introduces a novel method for analyzing 3D plant traits using a portable, affordable, and accurate tool, which has the potential to enhance plant breeding and agricultural practices.

**How to Use This Template:** The template details the sections that can be used in a manuscript. Note that each section has a corresponding style, which can be found in the “Styles” menu of Word. Sections that are not mandatory are listed as such. The section titles given are for articles. Review papers and other article types have a more flexible structure.

Remove this paragraph and start section numbering with 1. For any questions, please contact the editorial office of the journal or.

## 1. Introduction

The current global population of 7.7 billion people is expected to reach 9.8 billion by 2050 [1, 2], leading to increase in worldwide food demand. Moreover, due to the negative impact of climate change on the agriculture production and crop yields [3, 4, 5], new technological devices and approaches are required to be utilized, to precisely manage the limited resources and optimize crop yields by accurately monitor plants physiological and developmental responses to their environment [6]. Plant phenomics has exhibited significant promise in enhancing agricultural productivity through the systematic examination of plant physiological traits, such as growth, morphology, colors, size, dimensions, composition, and various other attributes.

The plant vegetative growth phase, particularly its leaves, plays a pivotal role in the growth and optimal productivity of plants. Leaves serve as the primary organs for photosynthesis, where carbohydrates are synthesized. Furthermore, leaves facilitate transpiration, the process of water evaporation from the plant into the atmosphere, enabling the upward movement of water and minerals from the soil. By quantifying leaf characteristics and monitoring their growth, valuable anatomical and physiological insights can be acquired. For instance, measuring parameters such as leaf length, area, and color distribution along the height profile can provide indicators of developmental stage, chlorophyll content, deficiencies and more. When combined with functional traits like transpiration rate and photosynthesis, a comprehensive understanding of the plant’s condition at any given moment can be obtained. Manual measurement of these attributes is time-consuming, inefficient, and potentially harmful to plants. Alternatively, utilizing specialized devices is costly and necessitates suitable infrastructure [7].

Remote sensing technology is an amerging domain in the field of percision agriculture, and plays a key element in its development [8]. Various technologies and equipments are available for monitoring crop growth and health, such as hyperspectral imaging (HSI), multispectral imaging (MSI) and Light Detection and Ranging (LiDAR) scanners [9].

LiDAR technology allows non-distractive, distant measurement through active laser beam illumination and facilitates the creation of a three-dimensional model of the subject being scanned, from which volumetric information, oragn-specific data and traits can be extracted. In recent years, LiDAR sensors received numerous attention in for field crop phenotyping and monitoring [10]. Some of the common lidar scanning facilities used are tripods, unmanned aerial vehicles and vehicles [11, 12, 13]. These technologies can be used to nondestructively extract accurate phenotypic information over a large spatial range and within a given time frame [14]. However, these technologies can be expensive and require complicated installation and specialized operations [15, 16]. The rapid technological advancement of smartphones together with low cost, made it possible to employ various sensors to collect plant data with high accuracy. For instance, the Apple iPhone 13 Pro smartphone is equipped with three high-quality cameras and a LiDAR scanner. Previous works have showed that the Apple iPhone 12 Pro LiDAR can be used to accurately model coastal cliffs [15] and was hypothesised that that 3D model can be created using Ipad Pro to extract specimen and habitats characteristics [18]. More advanced works have suggested automated process for trees stem diameter estimation using iPhone 13 Pro and an iPad Pro 2021 [19]. However, when compared with industrial scanner, iPad Pro 2021’s LiDAR scanner showed higher mean deviations of specific metrics when compared to real values, showing inpractical usage of small objects scanning [20]. Utilizing these iPhone capabilities offers a simpler and more affordable alternative to existing market products, thus making it an appealing choice for plant dynamics studies and parametric modelling.

The aim of this research was to test the hypothesis that accurate information, such as surface area, can be extracted from maize leaves (Zea mays) using the iPhone 13 Pro camera with LiDAR technology. This information would enable the calculation of diagnostic indicators for the plant while constructing a three-dimensional model that serves as a viable alternative to manual measurements. Thus, we generated three-dimensional models of cultivated maize that served as representative plant models for leaf measurements.

In this research we aimed to:

- Test the iPhone 13 Pro as a new handheld and affordable research tool, estimate its reliability and plant level data acquisition precision in the agriculture sector.
- Conduct weekly 3D scanning of maize plants that were grown under controlled and monitored conditions.
- Create 3D point cloud database of maze plants.
- Automate the process of plant organs segmentation of the point clouds to extract leaves and whole plant surface area and compare these traits to transpiration and growth data.

## 2. Materials and Methods

### 2.1 Experimental Design

The experimental and technical design of the study is presented in Fig. 1.

**Figure 1.**
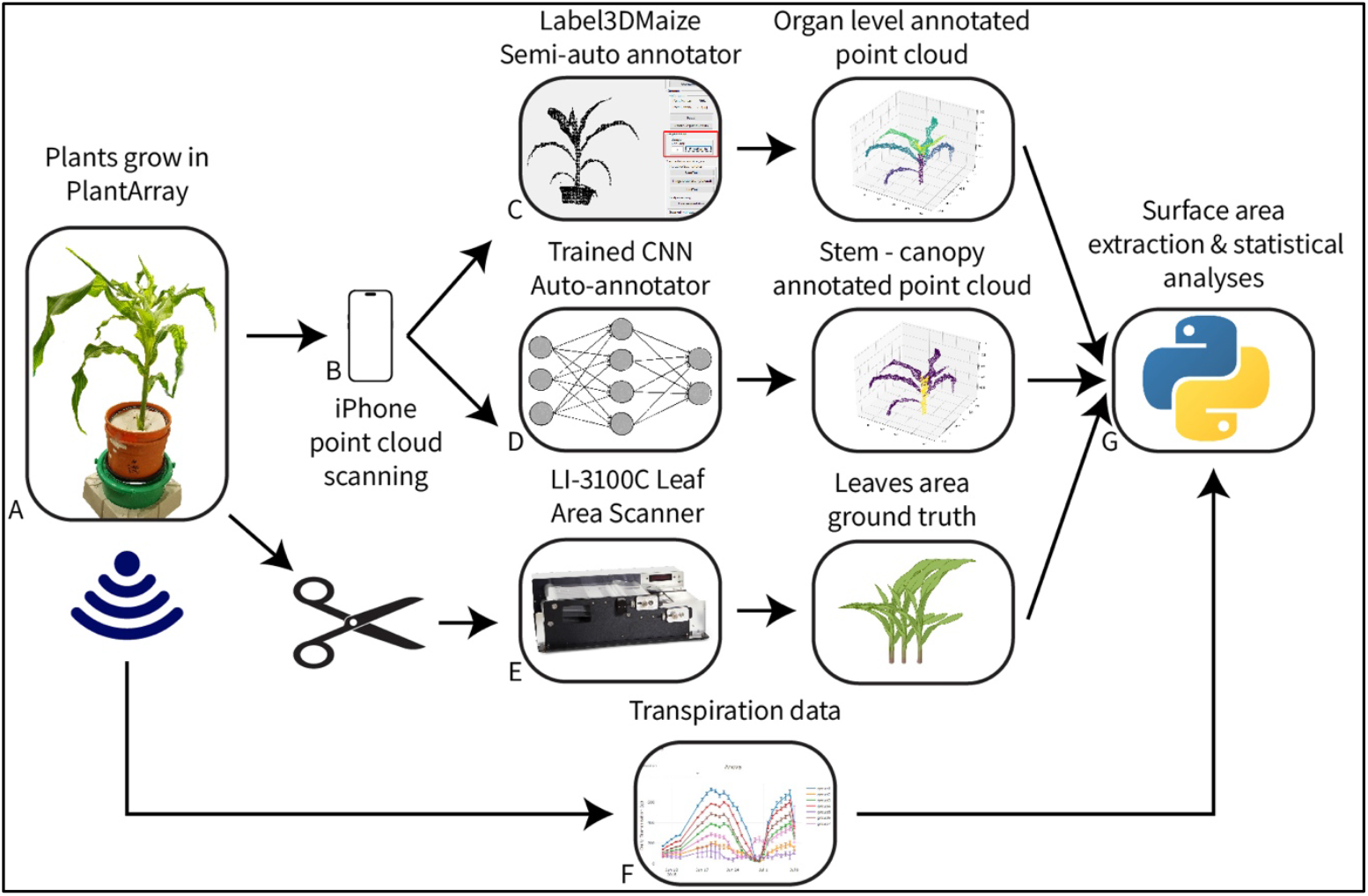
Overview of the experimental and technical design of the study. A) Maize plants were grown in PlantArray system, while transpiration data was constantly collected. B) Weekly 3D scanning of the plants conducted using iPhone to generate point clouds. The point clouds went through different annotation methods, including C) Label3DMaize software that was used to annotate each leaf separately in the point clouds, D) pre-trained CNN model to segment and annotate stem and canopy points separately. E) Leaves of scanned plants where cut and scanned using the LI-3100C scanner to produce ground truth data and corelated to F) the whole canopy transpiration rate. G) Python based algorithm was used to extract leaves surface area from the point clouds and conduct statistical analyses.

### 2.2 Technical testing

To test the accuracy of the Apple iPhone 13 Pro, 4 rectangular objects with flat surfaces were scanned using the ‘Polycam app’ (Polycam 3D Scanner App, LiDAR, 360 by Polycam Inc.) and their dimensions were measured with a measuring stick. The objects used in the test were matches box, iPhone box and wooden cubic boxes with different volumes. The dimensions of the objects scanned range from 6.5 × 1.72 × 11 cm up to 25 × 25 × 25 cm. Each object was scanned 3 times and the similarity to the real dimensions was evaluated. (Supplementary Table 4). The ‘Polycam app’ allows 2 types of scanning modes – LiDAR and Photo mode. We found that the LiDAR mode (utilizing the LiDAR sensor) is better for spatial scanning with when less accuracy needed, while the Photo mode suites objects scanning is much more precise (Therefore, all the data presented in this study utilize this option).

### 2.3 Data acuisition

Experiment took place at a growing room in the Faculty of Agriculture, Food and Environment. 15 Maize plants (hybrid line HMX59Y5832) were grown for 4 weeks under controlled environment using the Plant Array system. Average daily environmental conditions included 44.015 relative humidity (RH, %), 22.196 temperature (C), incoming flux radiation 475.818 (PPFD, 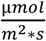). PlantArray (PlantArray System by Plant-DiTech LTD) is a high-throughput, multi-sensor physiological phenotyping gravimetric platform [21], enabeling the collection of individual discrete plants’ daily transpiration rates (Fig. 4A).

#### 2.3.1 Point cloud scans

Throughout the 4-weeks growing phase of the plants, starting from the 2nd week, weekly 3D scans were carried out individually for each plant (on the dates 2023-02-15, 2023-02-23 and 2023-02-28). Each plant was scanned by moving the iPhone in a semicircular motion along the vertical axis. This process ensured that all leaves were scanned both their upper and lower surfaces. Additionally, the iPhone was tilted around the plant, capturing data from 360 degrees along the horizontal axis (Fig. 2). After scanning is done, a ‘.xyz’ point cloud file was exported from the app for further analysis, explained next section.

**Figure 2.**
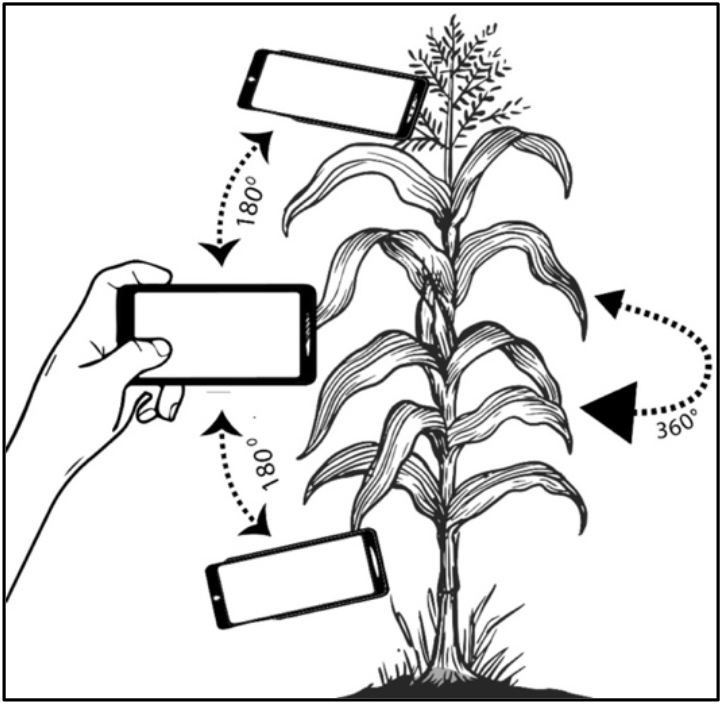
Demonstration of scanning process of a maize plant from all different angles to capture all details.

#### 2.3.2 Leaf area scans

At the end of the 4th week the plants were harvested, and each leaf went under the process of manual scanning using LI-3100C Area Meter Leaf Scanner. The scanner is a rugged benchtop instrument designed to quickly digitize the projected area, length, and width of leaves. It features two transparent conveyer belts that rotate to move leaves across a scanning bed. Each leaf was scanned 3 times (technical replicates) and the values were averaged to get the actual surface area of the leaves. This averaged value was then multiplied by a factor of two to align it with the point clouds’ estimated surface area.

### 2.4 Point cloud labeling and processing

Two approaces for point cloud segmentation and labeling were tested in this paper, semi-manual labeling using Label3Dmaize and auto segmentation using a pre-trained Convolutional Nneural Network (CNN) model.

#### 2.4.1 Manual labeling

Each point cloud was processed and labeled separately using the CloudcCompare and Label3Dmaize. CloudCompare was used to eliminate pot and surroundings points while keeping only the maize plant’s shoot and leaves (Fig. 3). Label3Dmaize [22] was then used to segment stem and individual leaves points separately (Fig. 5).

**Figure 3.**
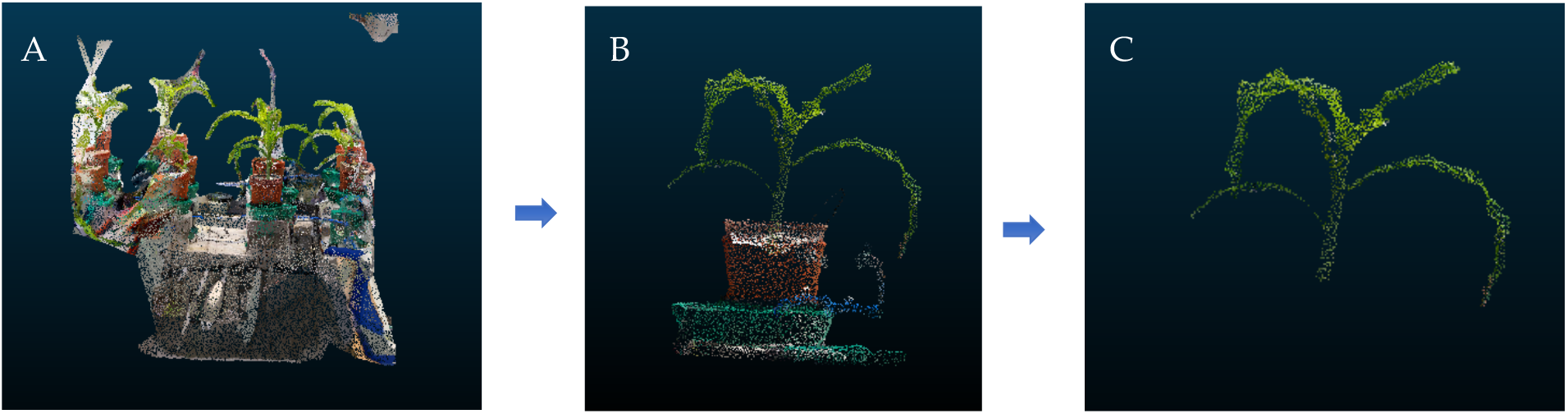
Point cloud editing process stages. Initial point cloud exported from the Polycam app. B) Point cloud with surroundings excluded. C) Final point cloud file of a maize plant with stem and leaves only.

**Figure 4.**
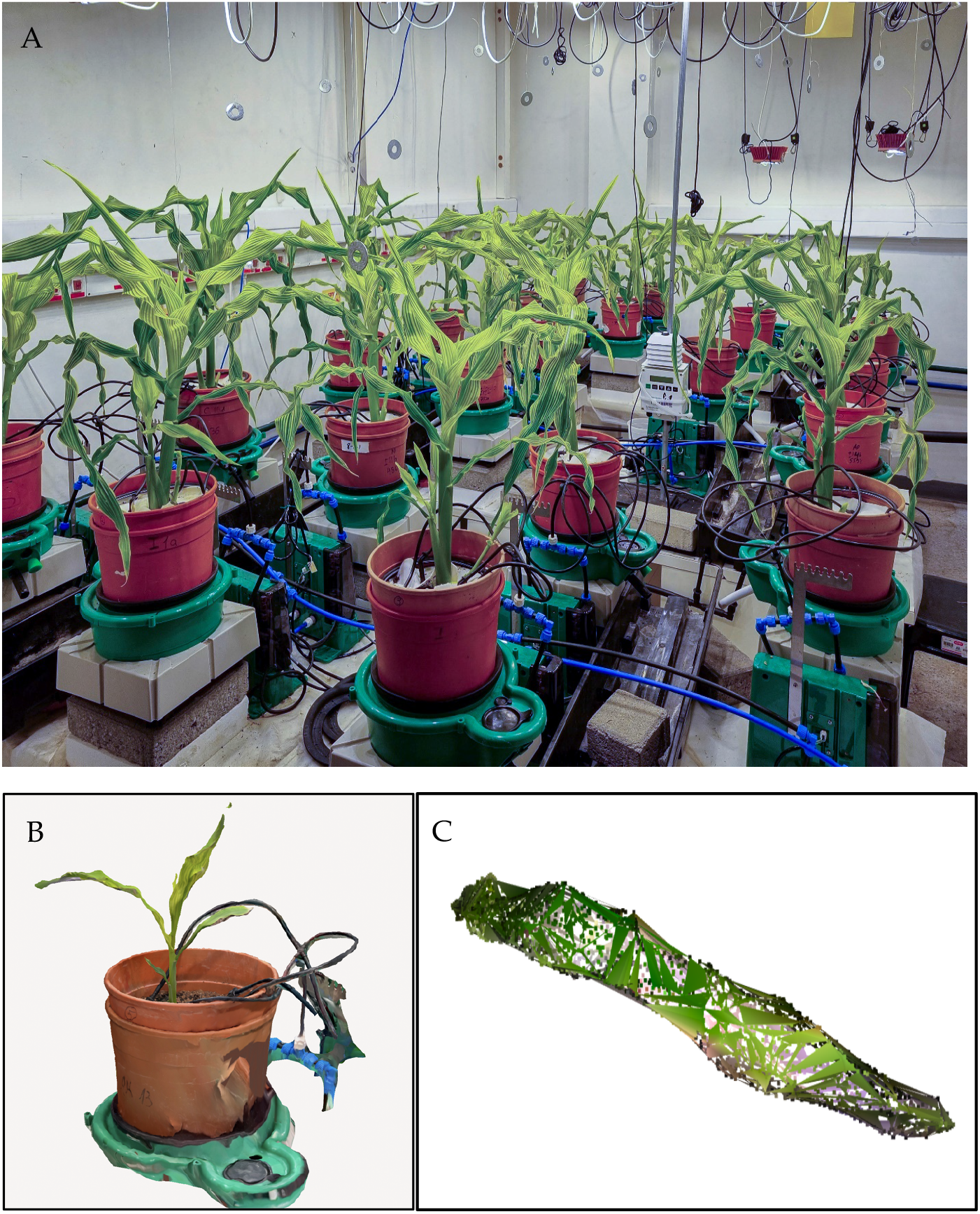
Evaluating iPhone accuracy for maize leaf 3D scanning and surface area measurement. A) 4 weeks old maize plants in a controlled growing room. Each plant is placed on top of separate lysimeter. B) Potted, whole plant scanning, presented as point cloud of a single maze, C) Surface constructed point cloud of a segmented leaf No.3, taken from B. The surface is constructed by triangles.

**Figure 5.**
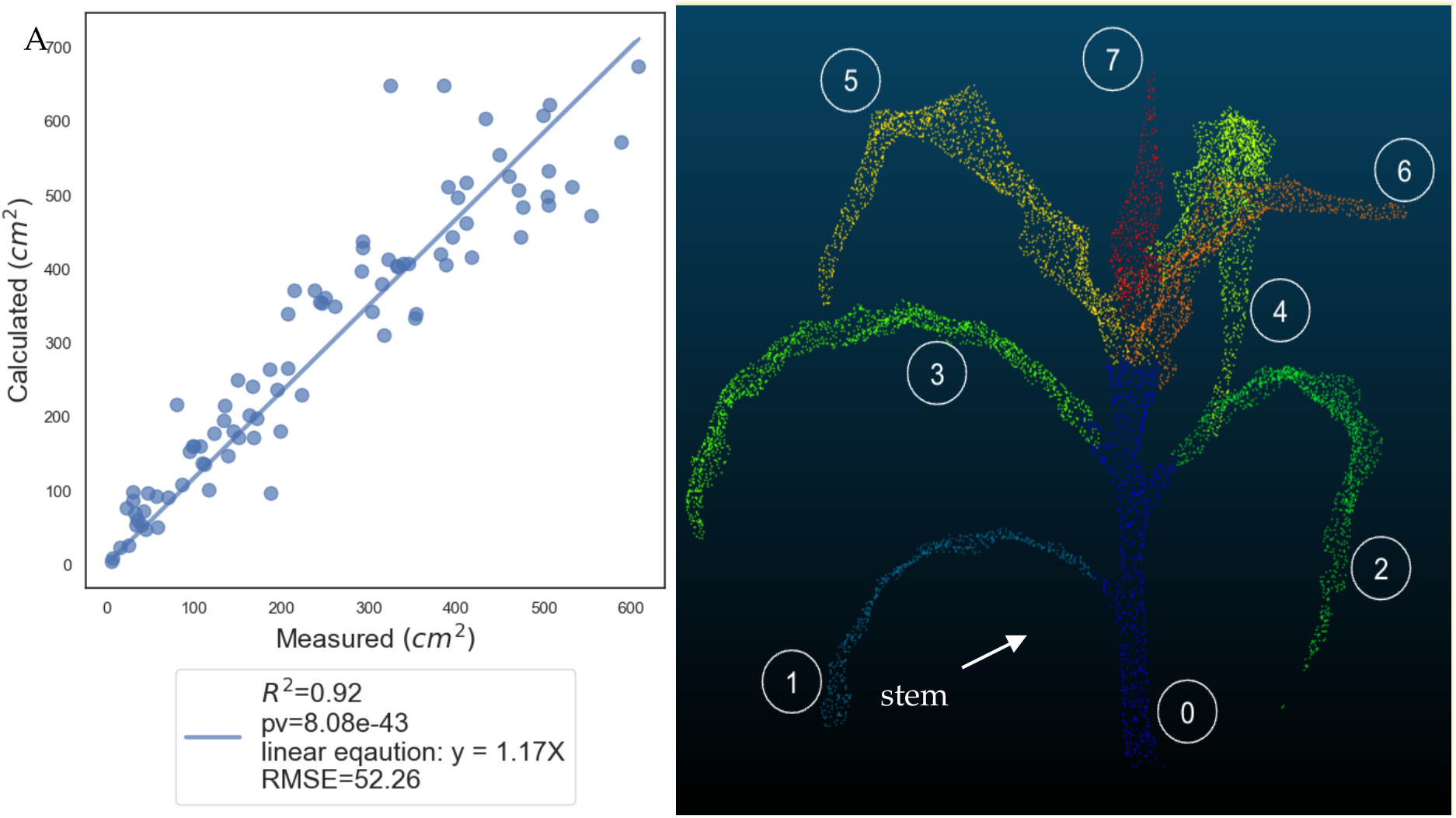
Comparative analysis of direct surface area measurements and computational extractions from point cloud data in maize leaves. A) Correlation between the manual surface area measurements and calculated ones, generated using the iPhone’s 3D scans. Shaded areas represent 95% confidence intervals of regression line. B) Annotated point cloud of a maize plant No.6 with an arrow pointing at the stem. Each leaf is colored separately and numbered according to its appearance along the stem (e.g. 1) from the bottom.

Label3Dmaize is an interactive corn plant point cloud segmentation and annotation software. The software enable the user to select the top and bottom stem points in the maize point cloud and adjusts the corresponding radius both via manual interaction. The median region growing method is then applied to achieve automatic segmentation and labelling of the maize stem and leaves. Suppose the connection area between the stem and leaves is not accurately annotated. In that case, the toolkit provides a fine segmentation function that allows correction of the misclassified points through simple manual interaction. After each point was labeled correctly, the Alpha Shapes algorithm was then utilized to generate surface reconstruction to further enable the extraction of the surface area of each leaf. The alpha shapes is a generalization of a convex hull [23], a computational geometric approach used for defining and analyzing complex shapes within point cloud data or sets of data points in Euclidean space. The algorithm takes its name from the parameter α, a parameter that influences the shape of the generated alpha shape. In our study we used α = 0.6. The basic idea behind the Alpha Shapes algorithm is to construct a series of nested shapes that represent the varying levels of connectedness between neighboring points. When the alpha parameter is decreased, the shapes gradually forming an increasingly complex boundary that encompasses the given data points. The Python package Open3D implements this algorithm to construct a triangle mesh from an unstructured point cloud while performing surface reconstruction. The surface is constructed by adjacent triangles, allowing the calculation of its area by using and summing the Heron’s formula for each triangle:

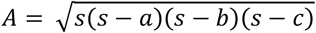

Where A is the triangle area; a, b and c represents the length of the triangle sides and s is computed as 0.5* (*a* + *b* + *c*). Mean and standard deviation metrics were computed for the measured and calculated surface areas of each leaf, with respect to their chronological appearance along the stem, across all plants (Table 2).

#### 2.4.2 CNN

Convolutional Neural Network (CNN) is a type of artificial neural network that is primarily used in image processing, computer vision, and other applications that involve spatial inputs. In this approach, we tested a trained CNN model for automatic stem and leaf segmentation for the maize point clouds (Ao et al., 2021) to distinguish between canopy and stem points, for the comparison of calculated whole canopy area with measured area. The model was designed to be executed on two separate platforms: Amazon Linux EC2 with Tesla K80 GPU (includeding CUDA9, a parallel computing platform, Python V3.6, Tensorflow V1.6 and NVIDIA driver version 384.98) and Amazon EC2 Windows Server 2022 to perform point cloud segmentation and stem extraction, respectively. For the segmentation, the first step of the model was down-sampling each point cloud into 2048 points, to suit the limitation of PointCNN the model was based on. The down-sampled point cloud was then tested by the model, resulting with initial segmentation of stem and leaf points. For the stem extraction, a 3D cylinder was fitted by the random sample consensus (RANSAC) algorithm around the stem segmented points (based on a given stem radius), considering that the maize stem grows vertically. The fitted cylinder position was then used to extract stem points in the initial point cloud, where points within the cylinders were labeled as stems and those left outside the cylinders as leaves.

### 2.5 Data analysis

The annotated point clouds were analysed in a few different ways, starting with the most detailed single leaf level to the whole canopy level.

#### 2.5.1 Single leaf level point cloud analysis

Initial testing included a comparison between manual measurement of a single leaf surface area (n=4, technical replicates) with the iPhone’s calculated surface area. For the accuracy assesment the %Error formula was used:

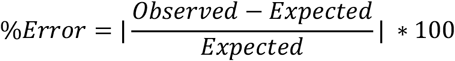

After confirming the accuracy level of the iPhone’s cameras and the surface reconstruction method, a linear regression was applied to compare the manually measured ground truth surface area of all leaves with the surface area derived from the point cloud data. In this section, 90 leaves (biological repetitions) from 15 different maize plants were tested altogeher. To assess the accuracy of the phenotypic trait extraction, the data was split into train and test sets (0.7 and 0.3, respectively). Both the coefficient of determination (R^2^) and the root mean squared error (RMSE) were used as quantitative metrics on the test set, calculated using the following equations:

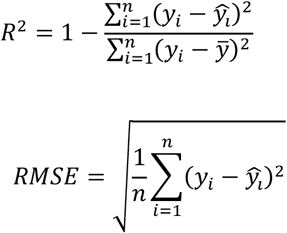

Where *y*_*i*_ is the actual value, *ŷ*_*j*_ is the predicted value, 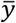 is the mean of actual values and *n* is number of observations.

#### 2.5.2 Simplified whole canopy level point cloud analysis

The method above required us to distinguish and separate the stem from the entire surface area, which proved to be a challenging task, therefore, the aim of this analysis was to test if canopy area can be estimated directly from the whole maize surface area.

Instead of segmenting the stem and canopy prior to comparison (as explained above), a regression analysis was conducted to establish a relationship factor between stem and total plant surface area. Whole plant surface area was calculated as:

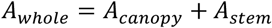

*A*_*stem*_ was computed as:

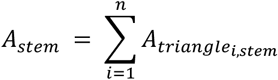

Where 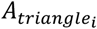 is the triangle area of the *n* total generated triangles during the surface reconstruction by the Alpha Shapes algorithm, using the stem labeled points. And

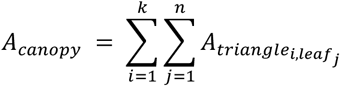

Where *k* is the number of leaves of each maize plant, and *n* is the total generated triangles during the surface reconstruction using the Alpha Shapes algorithm, using the leaf labeled points.

This approach enabled the prediction of canopy area based on the total plant total surface area (Materials, Table 1). For this analysis, data of 14 semi-automatically segmented and labeled maize point clouds was used. The resulted fited linear equation, stem area as a function of total area, was used to calculate the canopy area for each plant by subtracting the predicted stem area from the whole plant area.

#### 2.5.3 Simplified whole canopy level point cloud analysis using CNN model

Here, each point cloud underwent stem extraction three times using the trained CNN model, each time with a different radius parameter value (0.01, 0.02 and 0.03 m) for the 3D cylinder fitting, resulting with a total of 39 point clouds (2 of the 15 files could not be tested), with a varying degrees of successful segmentation achieved (Fig. 7, Materials - Table 2). Canopy area was calculated for each point cloud as based on the leaf segmented points. Linear rgression analysis was then conducted for each plant’s measured canopy area against its calculated canopy area, derived from the predicted canopy of the CNN model (represented by purple points in Figure 5, indicating leaf-labeled points).

#### 2.5.4 Comparing daily transpiration rate to canopy areas

Subsequently, we conducted a comparison between the daily transpiration rate of each plant, on a single date, and their respective calculated canopy areas derived from the different approaches mentioned earlier. The daily transpiration rate data was collected separately for each plant using the PlantArray system (methods, Fig. 4). Additionally, a One-Way ANOVA test was conducted to evaluate the differences in canopy area between the different approaches described above.

#### 2.5.5 Detecting growth and daily transpiration rate over time

Throughout the 4-weeks growing phase of the plants, the daily transpiration rate 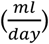 was recorded continuously and separately for each plant using the Plant Array system (Fig. 4A). A comparison between the transpiration rate and the whole surface, area on the same day of scanning (Fig. 11), was conducted. The decision was made to use the transpiration rate from ‘2023-02-26’ instead of ‘2023-02-28’, as on the latter date, the plants were harvested for leaf measurements and transpiration data collection was interrupted.

#### 2.5.6 Point cloud density

To investigate the potential impact of point cloud density on the %Error of surface area estimations, we conducted a comparison between point cloud density, calculated as:

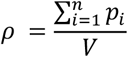

Where *n* is the number of points in the point cloud and *V* is the volume of the bounding box of the point cloud, which is the smallest box that contains all points. The dimensions f the box are determined by the minimum and maximum coordinates in the x, y, and z dimensions, as follows:

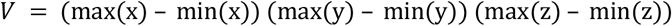

The mean %Error was determined as the average %Error between the calculated and measured surface areas for each leaf within a plant:

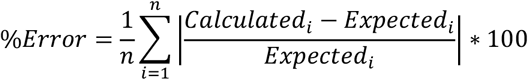

Where *n* is the number of leaves within each point cloud. Correlation between the density and %Error was tested, and both were compared separately with the number of frames 2D frames that were utilized by the Polycam app during the scanning process for point cloud construction through superposition. To explore the relationship between these measurements, linear regression was employed.

## 3. Results

### Testing the scanner with a rectangular objects and a single maize leaf

Comparing the dimensions of 4 rectangular boxes (n=3, technical replicates each), showed high mean similarity across all objects and sides of 90.26% (±8). Scanning accuracy is decreasing when object side length is below 5 cm (Supplemantery Table 4, Fig S1), similar to [25]. Manual measurements of a single leaf surface area averaged 34.625(±1.11;*cm*^2^, n=4, technical replicates), while the iPhone’s calculated surface area using an alpha parameter value of 0.6 was 71.022 (*cm*^2^). Comparing the two resulted in a percentage error of 2.5% (%Error formula, materials), thus confirming the accuracy of the scanning process and the effectiveness of the surface reconstruction method.

### Leaf level point cloud analysis

A comparison between the actual surface area and the estimated surface area of multiple leaves (n=90, biological repetitions) was conducted using linear regression analysis. The linear equation derived from the regression analysis of manual and 3D scan surface area measurements in maize leaves was Y = 1.17X with a *R*^2^ value of 0.92 and a significant correlation with a p-value less than 0.01. (Fig. 5). This analysis showed a deviation of 17% from the surface are measured by the leaf scanner. Average scanning time of a single plant was 2.38 (±0.33, min).stem

**Table 2.** Detailed representation of measurements areas multiplied by 2, as shown in the regression graph for assessing measurement repeatability. Column “Organ No.” represents each leaf’s chronological number as shown in Figure 3B. Columns “Plant No.6 iPhone Estimated Area” and “Plant No.6 LI-3100C Measured Area” show the data for plant shown in Figure 3B and “Plant No.6 Leaf %Error” represents the %error between these columns. Columns “Cross iPhone Estimated Mean” and “Cross iPhone Estimated Std” represent the mean and std values of calculated area of each leaf’s chronological number, across all plants (Example – row number 2 represents the data for leaf No.1, most bottom leaf of each plant). Columns “Cross LI-3100C Measured Mean” and “Cross LI-3100C Measured Std” represent the mean and std values of measured area of each leaf’s chronological number, across all plants.

| Organ | Organ No. | Plant No.6<br>iPhone<br>Estimated<br>Area<br>(cm<sup>2</sup>) | Plant No.6<br>LI-3100C<br>Measured<br>Area<br>(cm<sup>2</sup>) | Plant<br>No.6<br>Leaf<br>%Error | Cross<br>iPhone<br>Estimated<br>Mean<br>(cm<sup>2</sup>) | Cross<br>iPhone<br>Estimated<br>std<br>(cm<sup>2</sup>) | Cross<br>LI-3100C<br>Measured<br>Mean<br>(cm<sup>2</sup>) | Cross<br>LI-3100C<br>Measured<br>Std<br>(cm<sup>2</sup>) | Mean<br>Cross<br>%Error | N |
| --- | --- | --- | --- | --- | --- | --- | --- | --- | --- | --- |
| Stem | 0 | 338.52 | N/A | N/A | 247.75 | ±108.83 | N/A | N/A | N/A | 15 |
| Leaf | 1 | 171.83 | 150.58 | 14.11% | 152.33 | ±79.76 | 120.58 | ±67.5 | 36.79% | 15 |
| Leaf | 2 | 380.12 | 314.97 | 20.68% | 321.07 | ±137.84 | 251.02 | ±121.64 | 32.76% | 15 |
| Leaf | 3 | 499.00 | 504.10 | 1.01% | 474.26 | ±171.37 | 385.63 | ±154.09 | 32.33% | 15 |
| Leaf | 4 | 573.04 | 588.93 | 2.70% | 460.59 | ±175.96 | 412.78 | ±161.33 | 18.69% | 15 |
| Leaf | 5 | 526.58 | 460.15 | 14.44% | 334.44 | ±162.56 | 282.19 | ±142.63 | 29.75% | 15 |
| Leaf | 6 | 437.39 | 292.75 | 49.41% | 209.95 | ±146.63 | 149.42 | ±94.97 | 67.745 | 13 |
| Leaf | 7 | 136.76 | 111.76 | 22.36% | 105.65 | ±44.34 | 51.06 | ±34.08 | 170.16% | 7 |

### Simplified whole canopy level point cloud analysis

To distinguish and separate the stem area from the entire surface area, the relationship between the stem and total plant surface area was linearly modeled. This regression yielding a highly *R*^2^ value of 0.9 and a significant correlation with a p-value less than 0.01. The resulting linear equation from the regression analysis was Y = 0.123X – 22.928 (Fig. 6A), showing that the stem area is less than 12.3% of the plant total area at this stage of growth. Subsequently, utilizing the linear equation for stem area prediction, the surface area of the entire canopy for each plant was calculated by subtracting the predicted stem area from the whole plant area. Comparing the predicted surface area of the whole canopy with the measured surface area (calculated as the sum of the leaves surface areas for each plant) resulted in a *R*^2^ value of 0.83 and a significant correlation with a p-value less than 0.01 (Fig. 6 b).

**Figure 6.**
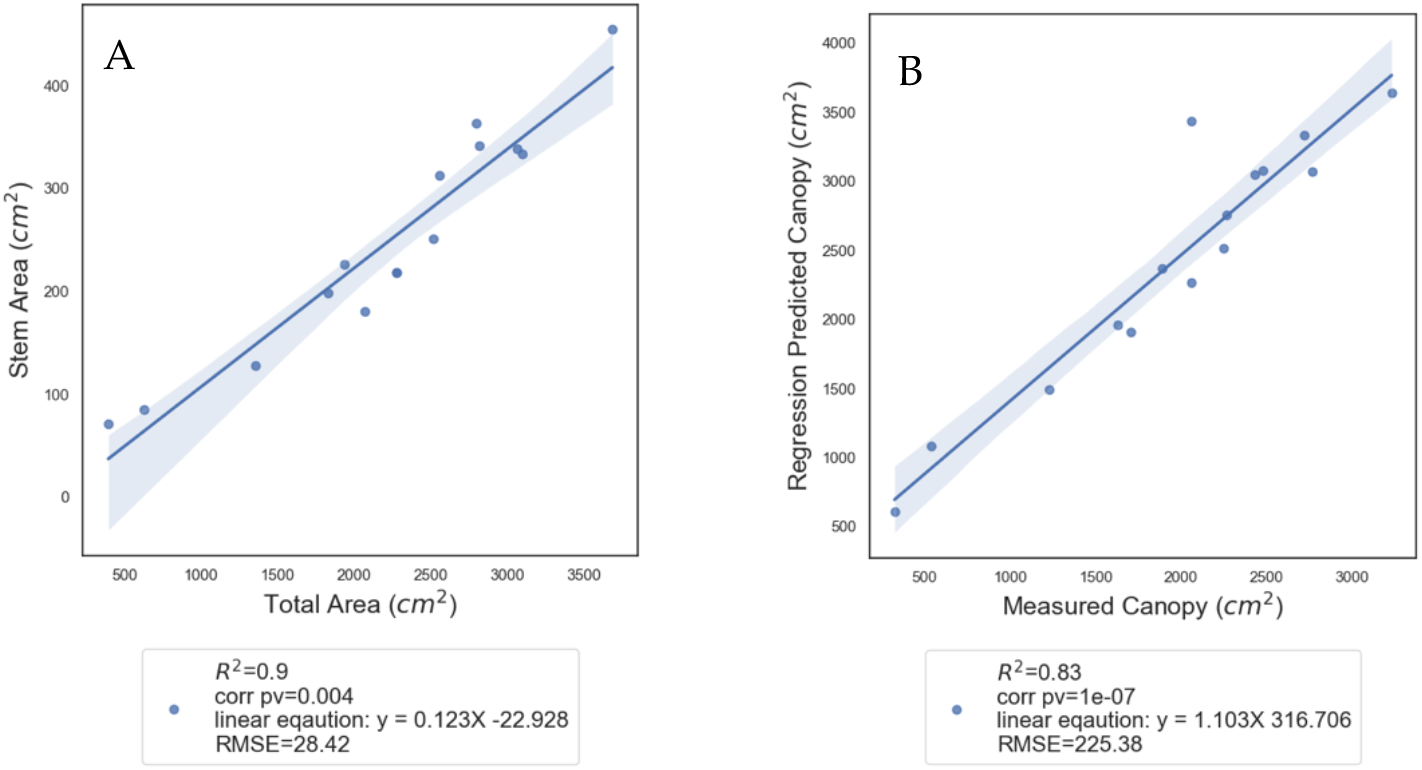
Predicting the whole canopy area of maize point cloud using stem area. Regression analysis was used: A) To predict stem area from plant total area. Using the resulted linear equation, the canopy area was predicted for each plant separately. B) The predicted canopy area was compared to the measured canopy area of each plant.

**Figure 7.**
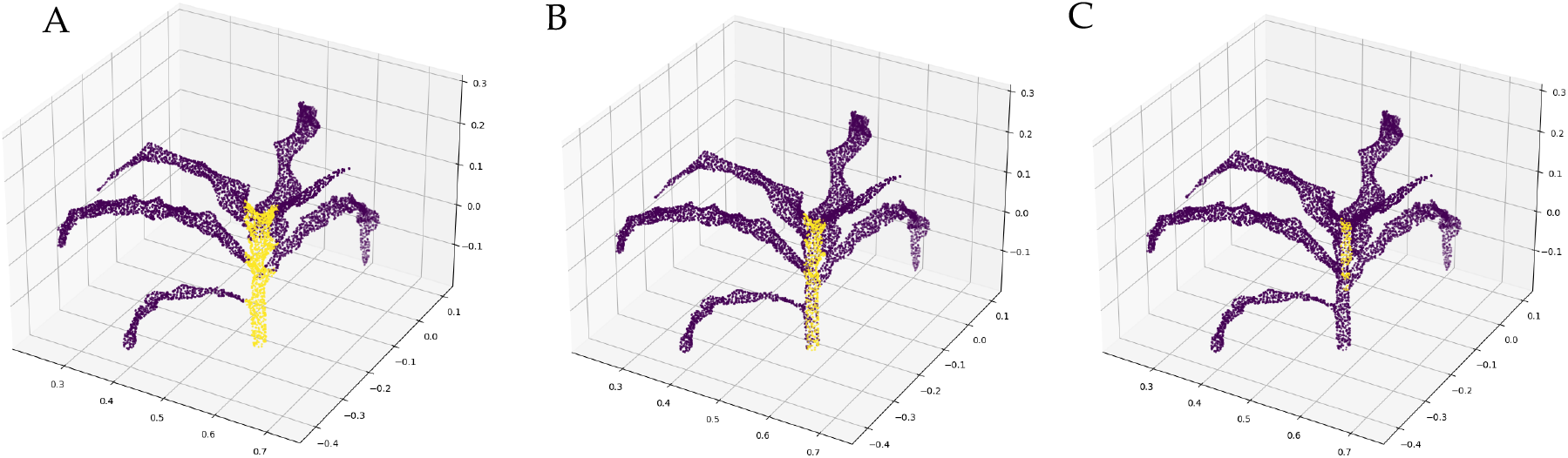
Impact of stem radius on maize canopy prediction accuracy. A) Fully egmented stem (yellow points) and leaves (purple points) using radius = 0.03m. B) Partially segmented stem points using radius = 0.02m, sporadic leaf labeled points in the stem. C) unsuccessful stem segmentation – almost all points (included the stem points) are labeled as leaf, using radius = 0.01m.

### Simplified whole canopy level point cloud analysis using CNN model

In this approach, we tested a trained CNN model for automatic stem and leaf segmentation for the maize point clouds (Ao et al., 2021) to distinguish between canopy and stem points, for the comparison of calculated whole canopy area with real area. Each point cloud underwent stem extraction three times, each time with a different radius parameter value (0.01, 0.02 and 0.03 m) for the 3D cylinder fitting, resulting with a total of 39 point clouds (2 of the 15 files could not be tested) with a varying degrees of successful segmentation achieved (Fig. 7, Materials - Table 2). Canopy area was calculated for each point cloud based on the leaf segmented points. Regression analysis was then conducted for each plant’s measured canopy area against its calculated canopy area, derived from the predicted canopy of the CNN model (represented by purple points in Figure 7, indicating leaf-labeled points). All three regression models exhibited a *R*^2^ values above 0.77 and a significant correlation with a p-value less than 0.01 (Fig. 8). On the graphs, green dots signify point clouds with successful stem segmentation, while red dots represent unclassified point clouds, containing only leaf-labeled points. In the latter case, the entire plant surface area was calculated as the canopy area (Materials – Table 2). The significant correlations observed, irrespective of stem and canopy separation, suggest that at this stage of plant growth, the stem area may not be relevant.

**Figure 8.**
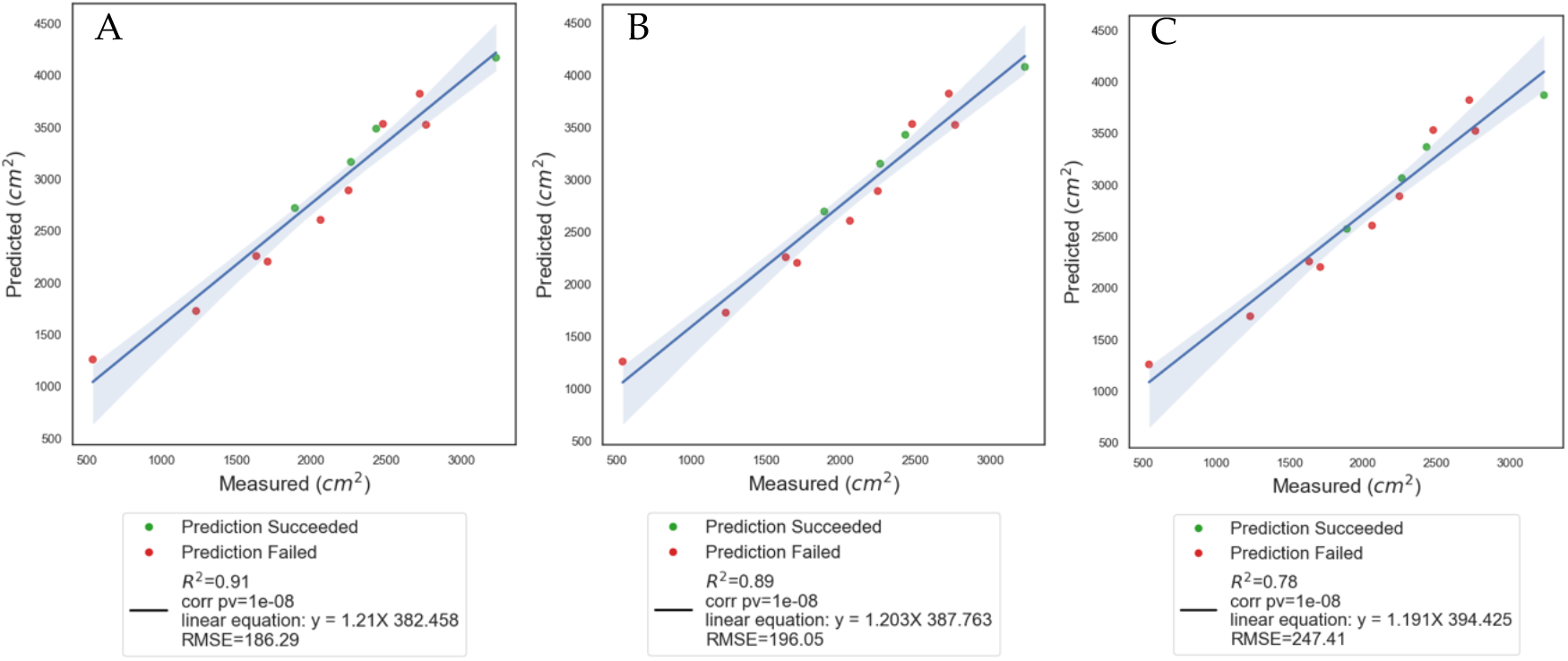
Comparison between canopy areas resulted by the CNN and the measured ones. Regression analysis for predicted canopy area by 3 radius parameter values: A) 0.01m, 0.02m and C) 0.03m respectively) against measured area. Green dots represent full successful stem segmentation point clouds, red dots represent point clouds with only leaf labeled points.

### Comparing daily transpiration rate to canopy areas

Correlations between daily transpiration rate and labeled canopy area, total area and regression factor canopy area resulted with a significant correlation with a p-value less than 0.01 and *R*^2^ values of 0.75, 0.75 and 0.81 respectively. Correlating the daily transpiration rate against canopy areas predicted using the CNN model, yielded significant correlation with a p-value less than 0.01 and *R*^2^ values of 0.81, 0.8, and 0.76 for testing with different radius parameter values of 0.01 (m), 0.02 (m) and 0.03 (m) respectively (Fig. 9).

**Figure 9.**
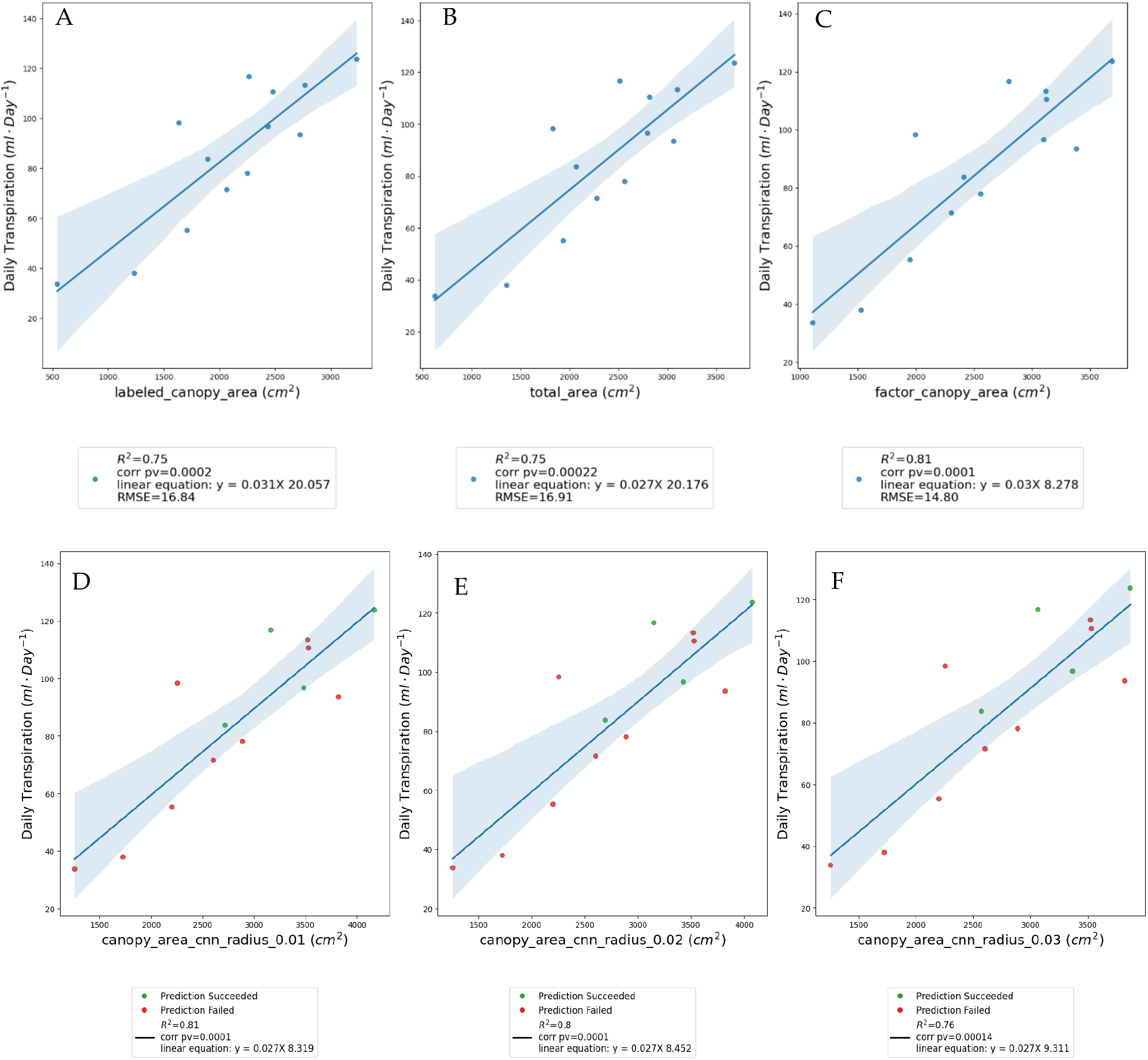
Comparing transpiration rate against canopy area predicted from different approaches. Daily transpiration linear regressions against: A) Semi-automatic labeled canopy area. B) Total area (stem + canopy). C) Predicted canopy by stem factor. D) CNN model radius = 0.01. E) CNN model radius = 0.02. F) CNN model radius = 0.03.

The One-Way ANOVA test results indicated no significant difference in canopy ar eacalculations with a p-value of 0.079 (Fig. 10). These results confirmed that the to tal plant area, which is the easiest to calculate and produce, can be utilized as an alternative to calculating the canopy area separately for transpiration predcition.

**Figure 10.**
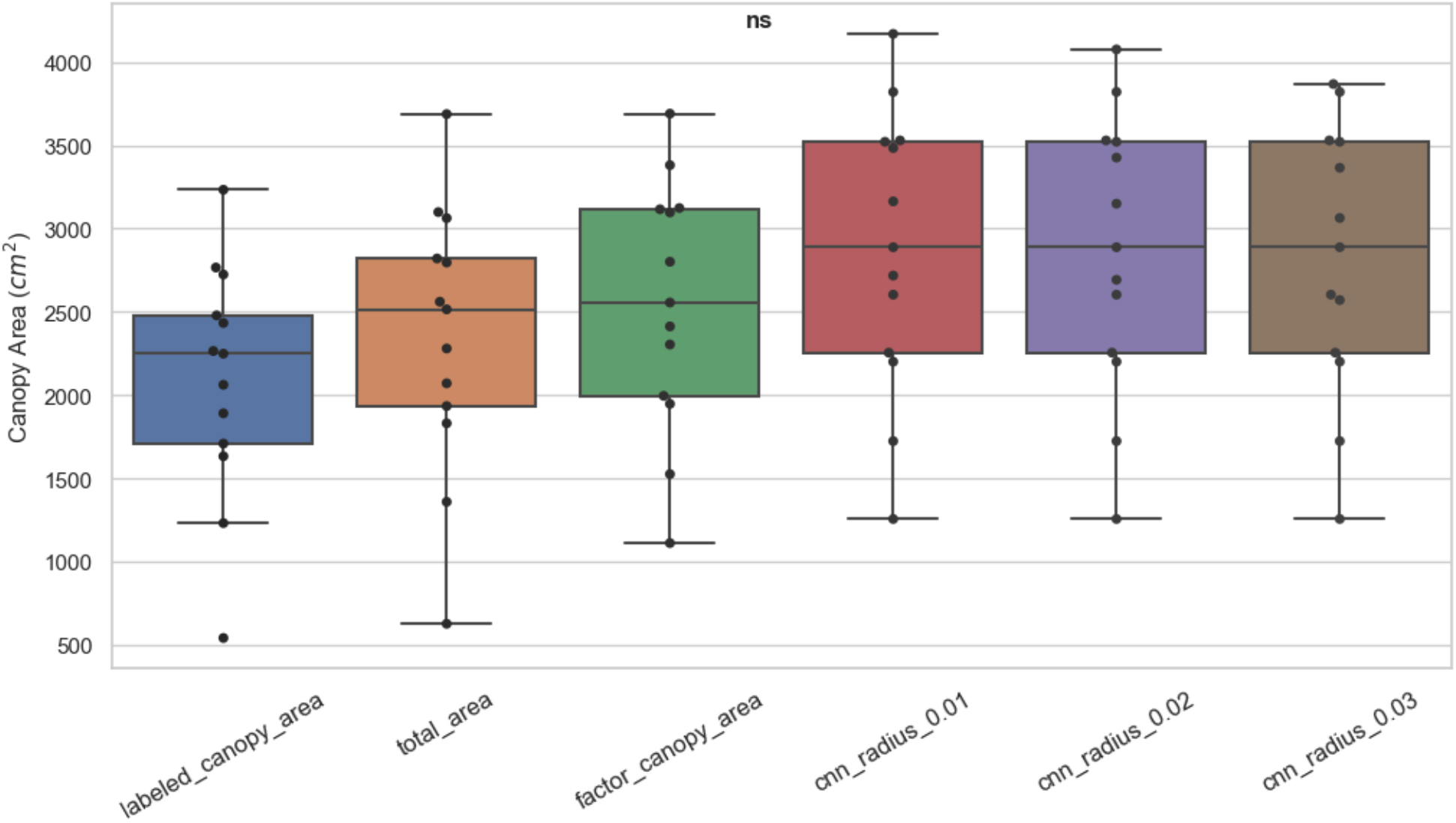
One-Way ANOVA test results for different transpiration predictors. Surface area distribution among different approaches. No significant difference between the approaches.

**Figure 11.**
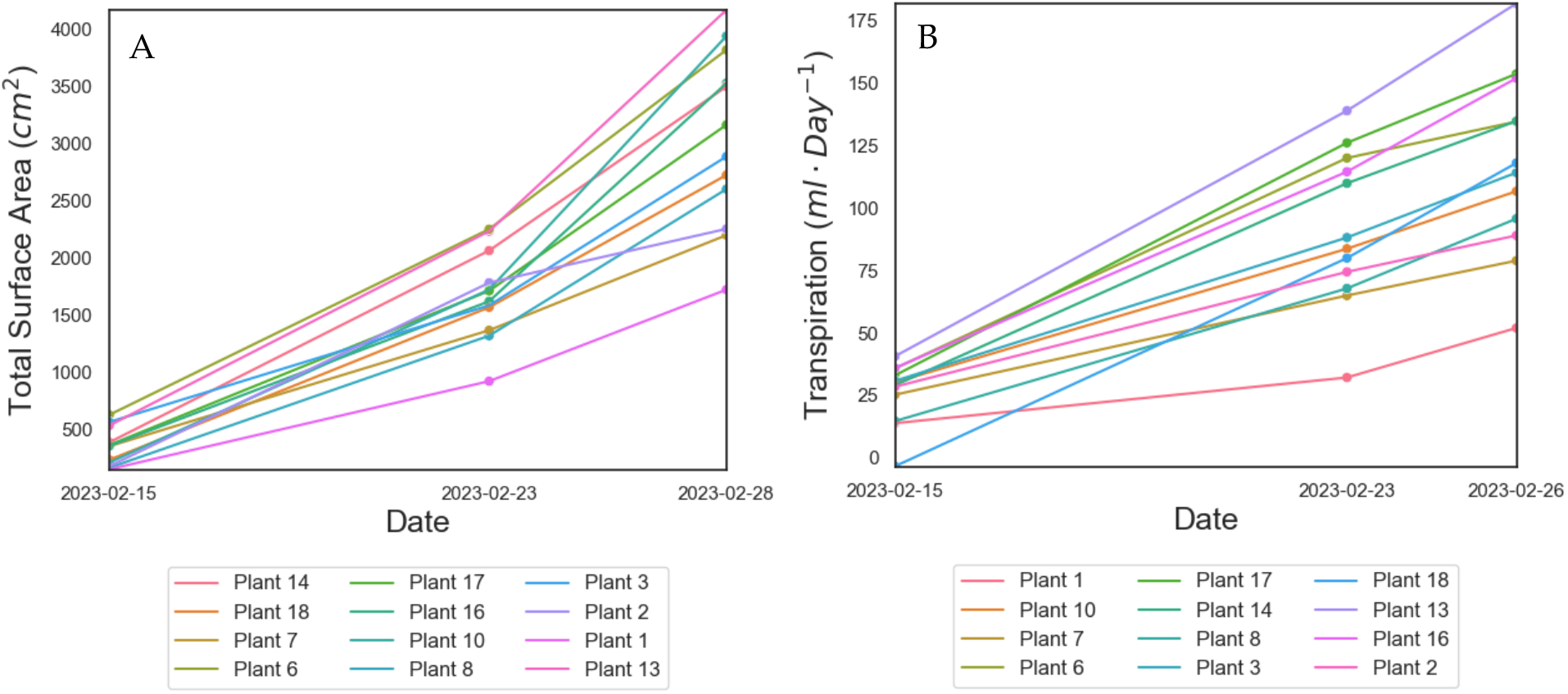
Change in plants’ surface area and daily transpiration rate. Change in plants’ growth over time, presented by A) change of total surface area (stem + canopy). B) Change in plants’ daily transpiration rates over time.

### Detecting growth and daily transpiration rate over time

A comparative analysis between growth and daily transpiraion rate using linear regression revealed a highly significant correlation with a p-value less than 0.01 and a *R*^2^ value of 0.83 (Fig. 12). This strong correlation between the plant’s growth rate, determined from the whole plant surface area, and the daily transpiration rate demonstrates the interrelation between plant development and water loss through transpiration.

**Figure 12.**
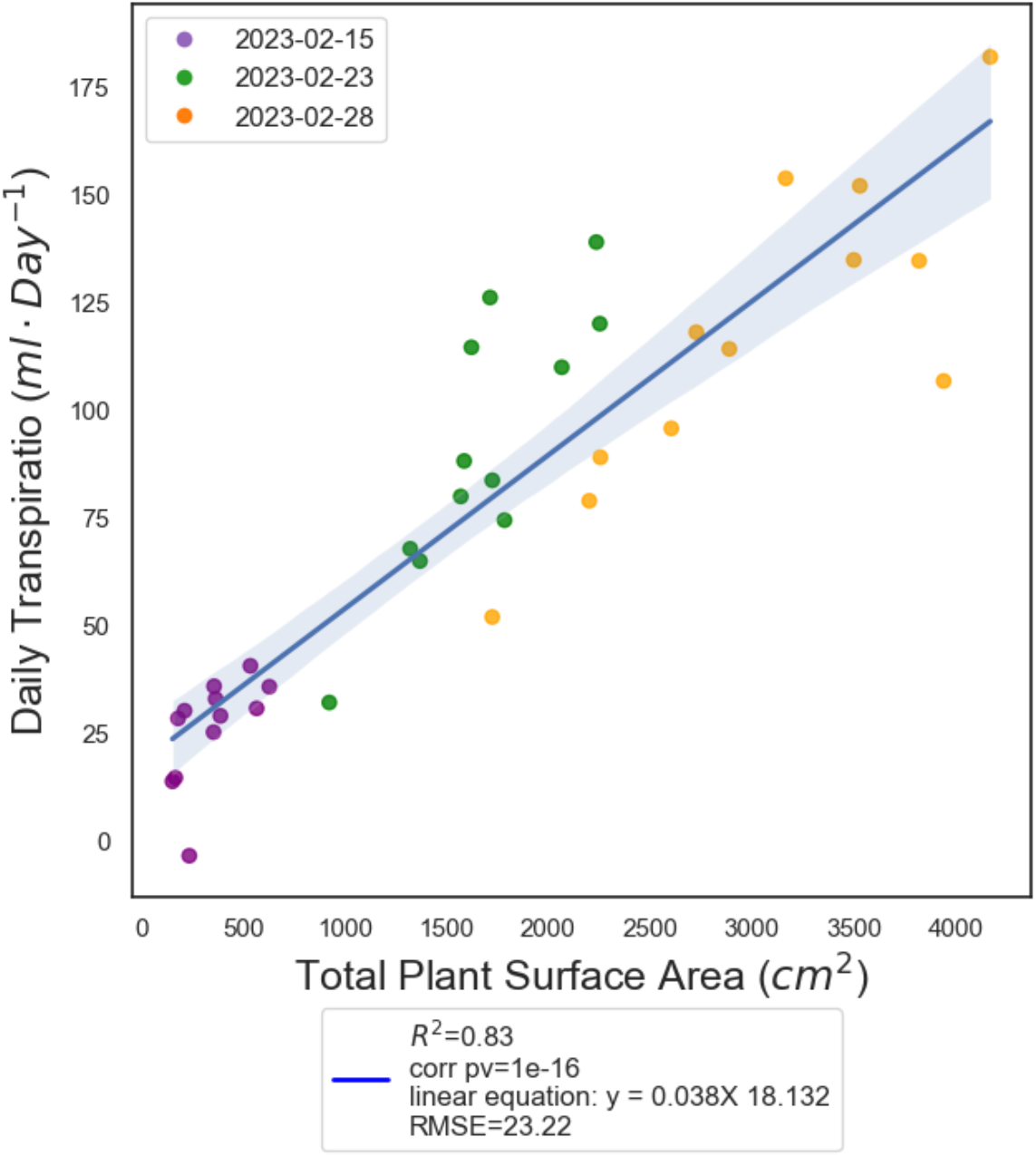
Plants’ daily transpiration rate vs. total surface area. Data for over the period of three weeks, each date is colored separately. Date ‘2023-02-28’ represents 3D scan from the ‘2023-02-28’ and transpiration data collected from ‘2023-02-28’.

### Point cloud density

Comparison between the density, %Error and number of frames, resulted in low *R*^2^ values and non-significant p-values (Fig. 13). These findings suggest that variations in the number of frames per file or the point density do not significantly impact the deviation in leaf area calculations. Therefore, it appears that altering the point cloud density within the tested range does not substantially affect the accuracy of surface area calculations for the leaves.

**Figure 13.**
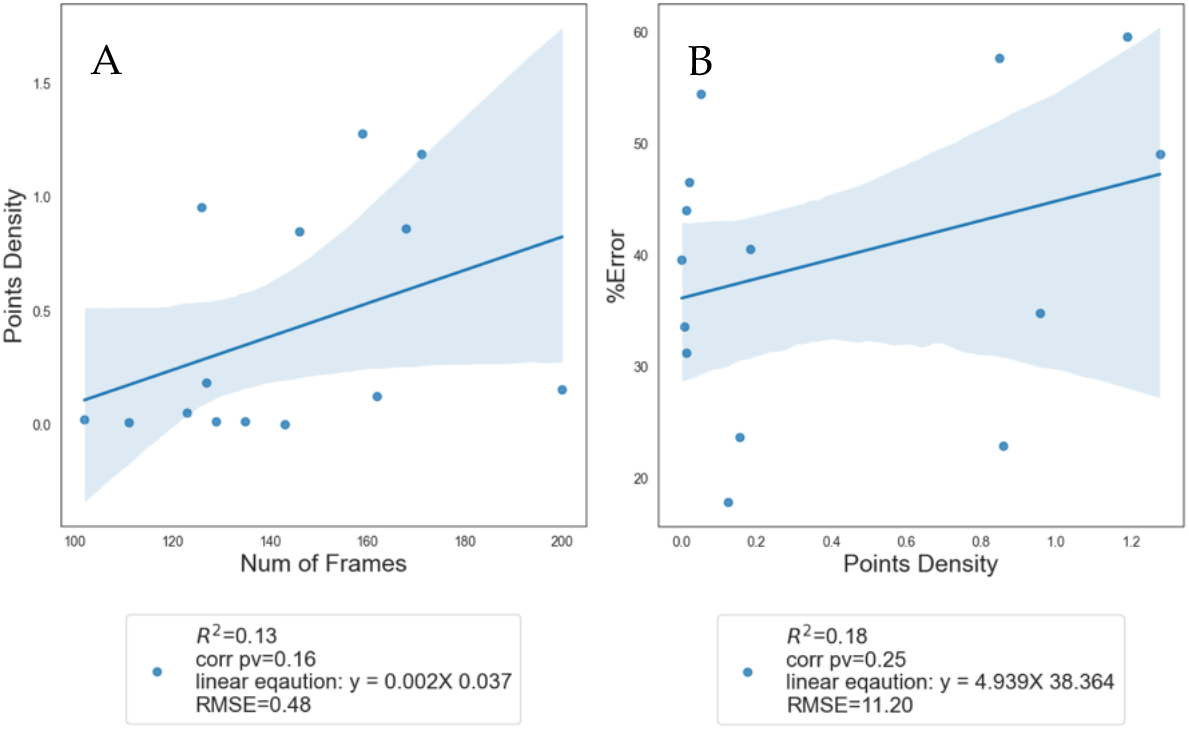
Nonsignificant comparisons of leaf area %error and point cloud density. Number of frames per point cloud vs. points density generated by the Polycam app point cloud creation. A) Positive linear relationship between point density vs. number of frames. B) Positive linear relationship between %error and point density.

## 4. Discussion

In this study, we investigated various aspects of plant phenotyping using 3D point cloud data generated using iPhone 13 Pro. Our findings shed light on the potential of this approach for tracking plant growth and whole canopy surface area computations, while using an accurate and more affordable and an easy to use measuring tool than those used in previous works [24, 25, 26, 27].

The iPhone’s factory calibration has been proven to be very accurate (Fig. 4, Fig. 5A), saving the user a tedious calibration process, which serves as a significant advantage in general and in particular to scientific goal, as a measuring tool for object sizes.

The accuracy of surface area computations derived from point cloud data revealed that individual maize leaf trait extraction using manually labeled point clouds resulted with a high *R*^2^ value of 0.92, when compared with reference measured area (Fig. 5A), similar to the results showed in [24, 25] when tested more advanced segmentations methods. We suggest that the 17% deviation between the calculated and measured surface area is not an error but reflect the true surface area of the leaf. This is due to the curved surface area the maize leaf, which results in larger area than the projected surface area derived from the leaf area scanner. Since the leaf area scanner only performs 2D scanning (projected area), it cannot capture the full 3D surface, as the iPhone scan does.

Next, we proposed a way for computing the whole canopy surface area by constructing a stem-total area ratio to avoid stem and canopy segmentation prior to comparison. Comparing the computed canopy area to the reference area resulted in *R*^2^ value of 0.83. We demonstrated that the computed whole canopy surface area of maize plants using CNN-based segmentation approach exhibited strong correlations *R*^2^ ≥ 0.78 () with the measured surface area (Fig. 8), though only 27% maize point clouds were segmented successfully, compared to [24], where 100% of stem instances were successfully extracted. Unsuccessful stem extraction in the second part of the model could arise from various stem angles in the maize plants, as the model assumes that stem grows vertically, hence the angle between the stem and z-axis should be less than 10 degrees.

Another significant scientific advantage of the iPhone scanning approach is that this manually-operated tool effectively generates a comprehensive 3D structure of the absolute plant canopy area. Contrasting with high-end machines that traverse over plants to produce only the top canopy’s 3D projected area, the iPhone not only demonstrates a robust correlation but also a highly accurate approximation between Y and X (Y= 1.17X, Fig. 5). Meanwhile, top canopy scanners, although establish a decent correlation between their 3D scans of the top canopy and the observed scans, in reality introduce an error margin of ∼ 70% in measuring absolute values of the leaf area (Y=0.22X), as detailed in [16]. Hence, it is advisable to exercise caution when interpreting biological and agronomic traits related to actual leaf area, such as transpiration, especially when relying on measurements from above-canopy machines. Our, approach revealed that the whole canopy area values computed using the different methods were highly corelated to the whole plant transpiration. Moreover, we found no significant difference between our different analyzing methods (Fig. 10), suggesting that using the whole plant surface area could be used for transpiration prediction in maize grown under well-irrigated condition.

Then we investigated the relationship between daily transpiration rate and the plant total surface area. Remarkably, we observed a significant correlation (p-value < 0.01) and a high *R*^2^ value of 0.83, implying that transpiration is closely linked to plant surface area (Fig. 12). This finding aligns with existing knowledge, as transpiration is a fundamental process in plants, regulating water uptake and nutrient transport and is directly affected by canopy area [28]. The resulting model, represented by the equation Y = 0.038X + 18.132, indicates that even in the absence of total plant surface area (X = 0) and without stomata to transpire from, there exists a basal level of transpiration rate at 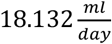. This observation implies a certain degree of inherent variability, potentially originating from factors such as ground surface evaporation. Although the ground surface may be covered, it is not entirely sealed to allow for roots’ respiration (as illustrated in Fig. 4A, 23). Additionally, other sources of data variability during collection may contribute to this observation.

Our findings suggests that the 3D point cloud data, acquired through the iPhone, can be effectively utilized to estimate the surface area of maize leaves. The high R2 and P values, indicate the reliability of the employed methods and highlight the potential for these techniques to be adopted in plant phenotyping studies. We also demonstrated that separating stem area for surface are calculation may not be relevant at this stage of the plant’s growth, as the ratio of the stem to the rest of the plant is 12.3% (Fig. 6A) and leaves covering the stem surface are also transpiring (Fig. 9B).

Lastly, we explored the effect of point cloud density on surface area calculations. Surprisingly, we found that altering the number of frames per file or the point density did not significantly impact the deviation in leaf area calculations (Fig. 13), as opposed to [27] that showed that with decreasing resolution the error is increasing. This suggests that within the tested range, variations in point cloud densities have minimal effect on the accuracy of surface area calculations. Therefore, researchers can potentially adopt different point cloud densities based on their specific experimental needs without compromising the reliability of the results, thus saving more time while performing scans and reduce storage usage.

Our study also demonstrated the feasibility of using smartphone-based technologies for plant phenotyping. The iPhone 13 Pro’s advanced features, such as three high-quality cameras, facilitated the creation of accurate 3D models of maize plants. This approach offered a more affordable and accessible alternative to dedicated devices available on the market, making it a viable option for plant researchers. Additionally, the successful utilization of smartphone-based technology highlights the potential for wider adoption of similar approaches in plant sciences, opening new avenues for understanding plant physiology and enhancing crop productivity.

Taking into consideration that our current approach would be complicated to up-scale, future plans including usage of robotic arm in addition to utilizing NeRF technology to be able to fully automate the process without the dependent of an app.

Considering the smartphone’s price, accuracy, ease of use, and self-calibration, it presents a stark contrast to the costly, high-maintenance, and infrastructure-intensive commercial “over canopy” scanners. These more expensive systems are often feasible only in large, well-funded greenhouses. The accessibility and practicality of the iPhone scanner make it an ideal tool for any research student, scientist, or field practitioner seeking reliable and efficient data collection methods.

## Supplementary Materials

**Table 1.** Maize point clouds data for establishing stem-total area relationship, consisting on the canopy area, stem area and total aera (canopy+stem).

| plant_num | canopy_area<br>($cm^2$) | stem_area<br>($cm^2$) | total_area<br>($cm^2$) |
| --- | --- | --- | --- |
| 1 | 1231.693 | 127.679 | 1359.372 |
| 2 | 1633.31 | 197.917 | 1831.227 |
| 3 | 2249.073 | 312.06 | 2561.133 |
| 6 | 2724.709 | 338.519 | 3063.228 |
| 7 | 1709.142 | 225.541 | 1934.683 |
| 8 | 2062.331 | 218.019 | 2280.35 |
| 9 | 2766.653 | 332.848 | 3099.501 |
| 10 | 2062.331 | 218.019 | 2280.35 |
| 11 | 541.852 | 85.17 | 627.022 |
| 13 | 3233.65 | 454.37 | 3688.02 |
| 14 | 2433.586 | 363.048 | 2796.634 |
| 15 | 325.171 | 70.996 | 396.167 |
| 16 | 2479.284 | 341.62 | 2820.904 |
| 17 | 2265.225 | 250.311 | 2515.536 |
| 18 | 1891.095 | 180.178 | 2071.273 |

**Table 2.** CNN model output and reference canopy area (measured_canopy_area) for each plant and radius parameter used. If stem segmentation was successful (model_log = ‘succeeded’) then canopy and stem area were estimated (under calculated_area). If stem segmentation did not succeed (model_log = ‘failed’) then only canopy area was estimated.

| plant_num | organ_type | calculated_area<br>($cm^2$) | Radius (cm<br>) | measured_canopy_area<br>(cm) | model_log |
| --- | --- | --- | --- | --- | --- |
| 18 | leaf | 2718.277 | 0.01 | 1891.095 | succeeded |
| 18 | stem | 116.43 | 0.01 | 1891.095 | succeeded |
| 18 | leaf | 2692.914 | 0.02 | 1891.095 | succeeded |
| 18 | stem | 294.056 | 0.02 | 1891.095 | succeeded |
| 18 | leaf | 2571.251 | 0.03 | 1891.095 | succeeded |
| 18 | stem | 413.831 | 0.03 | 1891.095 | succeeded |
| 9 | leaf | 3521.663 | 0.01 | 2766.653 | failed |
| 9 | leaf | 3521.663 | 0.02 | 2766.653 | failed |
| 9 | leaf | 3521.663 | 0.03 | 2766.653 | failed |
| 11 | leaf | 1256.367 | 0.01 | 541.852 | failed |
| 11 | leaf | 1256.367 | 0.02 | 541.852 | failed |
| 11 | leaf | 1256.367 | 0.03 | 541.852 | failed |
| 7 | leaf | 2201.025 | 0.01 | 1709.142 | failed |
| 7 | leaf | 2201.025 | 0.02 | 1709.142 | failed |
| 7 | leaf | 2201.025 | 0.03 | 1709.142 | failed |
| 16 | leaf | 3529.927 | 0.01 | 2479.284 | failed |
| 16 | leaf | 3529.927 | 0.02 | 2479.284 | failed |
| 16 | leaf | 3529.927 | 0.03 | 2479.284 | failed |
| 6 | leaf | 3820.954 | 0.01 | 2724.709 | failed |
| 6 | leaf | 3820.954 | 0.02 | 2724.709 | failed |
| 6 | leaf | 3820.954 | 0.03 | 2724.709 | failed |
| 17 | leaf | 3163.741 | 0.01 | 2265.225 | succeeded |
| 17 | stem | 65.975 | 0.01 | 2265.225 | succeeded |
| 17 | leaf | 3149.995 | 0.02 | 2265.225 | succeeded |
| 17 | stem | 240.616 | 0.02 | 2265.225 | succeeded |
| 17 | leaf | 3064.108 | 0.03 | 2265.225 | succeeded |
| 17 | stem | 385.617 | 0.03 | 2265.225 | succeeded |
| 1 | leaf | 1724.155 | 0.01 | 1231.693 | failed |
| 1 | leaf | 1724.155 | 0.02 | 1231.693 | failed |
| 1 | leaf | 1724.155 | 0.03 | 1231.693 | failed |
| 8 | leaf | 2603.65 | 0.01 | 2062.331 | failed |
| 8 | leaf | 2603.65 | 0.02 | 2062.331 | failed |
| 8 | leaf | 2603.65 | 0.03 | 2062.331 | failed |
| 3 | leaf | 2888.46 | 0.01 | 2249.073 | failed |
| 3 | leaf | 2888.46 | 0.02 | 2249.073 | failed |
| 3 | leaf | 2888.46 | 0.03 | 2249.073 | failed |
| 2 | leaf | 2255.337 | 0.01 | 1633.31 | failed |
| 2 | leaf | 2255.337 | 0.02 | 1633.31 | failed |
| 2 | leaf | 2255.337 | 0.03 | 1633.31 | failed |
| 13 | leaf | 4169.738 | 0.01 | 3233.65 | succeeded |
| 13 | stem | 156.984 | 0.01 | 3233.65 | succeeded |
| 13 | leaf | 4076.57 | 0.02 | 3233.65 | succeeded |
| 13 | stem | 460.058 | 0.02 | 3233.65 | succeeded |
| 13 | leaf | 3868.557 | 0.03 | 3233.65 | succeeded |
| 13 | stem | 648.728 | 0.03 | 3233.65 | succeeded |
| 14 | leaf | 3484.345 | 0.01 | 2433.586 | succeeded |
| 14 | stem | 167.026 | 0.01 | 2433.586 | succeeded |
| 14 | leaf | 3426.635 | 0.02 | 2433.586 | succeeded |
| 14 | stem | 337.275 | 0.02 | 2433.586 | succeeded |
| 14 | leaf | 3365.779 | 0.03 | 2433.586 | succeeded |
| 14 | stem | 476.797 | 0.03 | 2433.586 | succeeded |

**Table 3.**
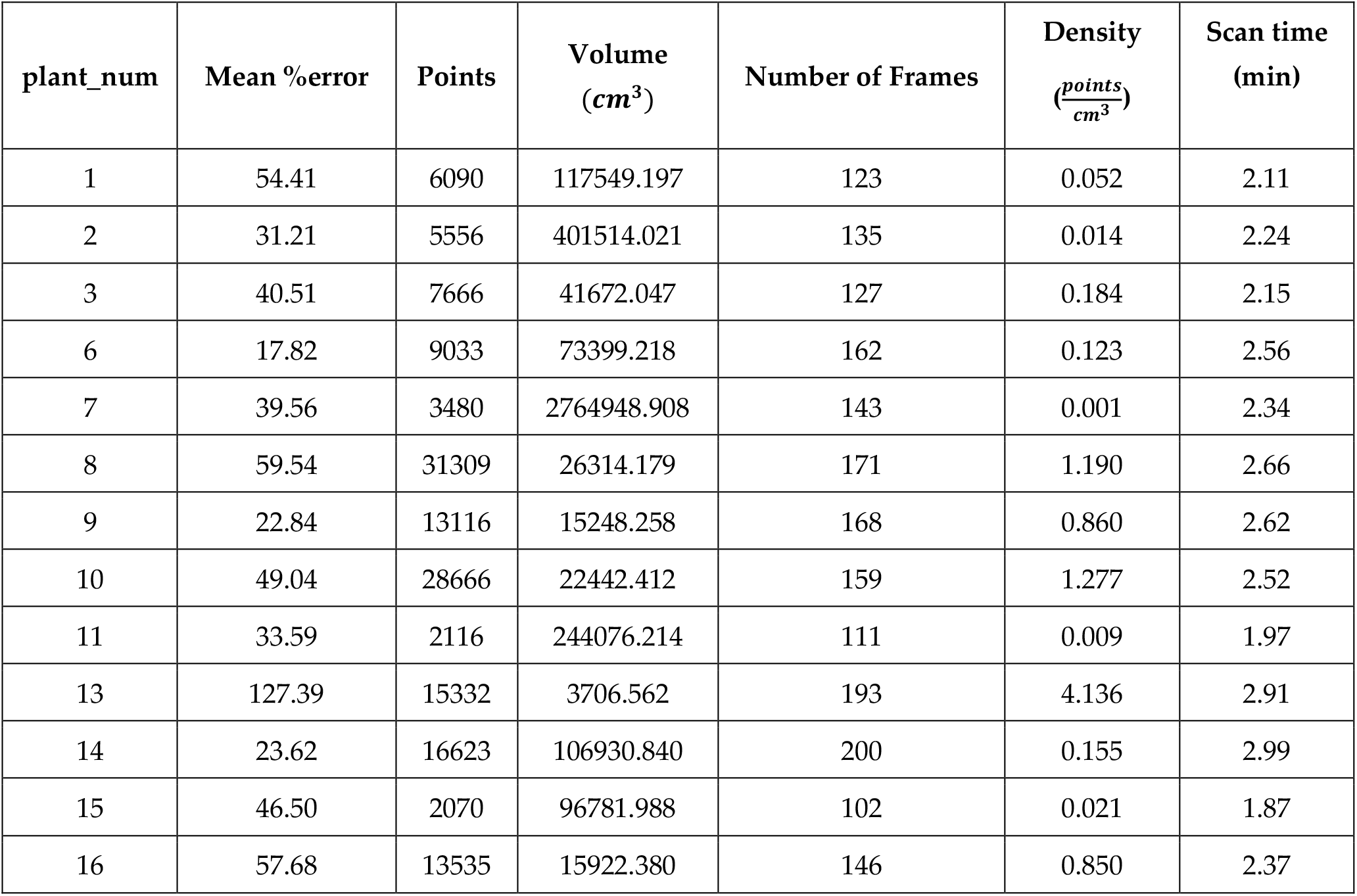
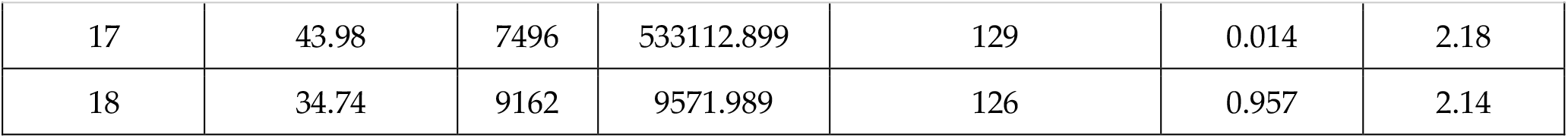
Point cloud densities including mean %error of leaves area for each plant, number of points and the volume of each point cloud, number of 2D frames used to generate the point cloud via Polycam app and point density.

| plant_num | Mean %error | Points | Volume<br>($cm^3$) | Number of Frames | Density<br>($\frac{points}{cm^3}$) | Scan time<br>(min) |
| --- | --- | --- | --- | --- | --- | --- |
| 1 | 54.41 | 6090 | 117549.197 | 123 | 0.052 | 2.11 |
| 2 | 31.21 | 5556 | 401514.021 | 135 | 0.014 | 2.24 |
| 3 | 40.51 | 7666 | 41672.047 | 127 | 0.184 | 2.15 |
| 6 | 17.82 | 9033 | 73399.218 | 162 | 0.123 | 2.56 |
| 7 | 39.56 | 3480 | 2764948.908 | 143 | 0.001 | 2.34 |
| 8 | 59.54 | 31309 | 26314.179 | 171 | 1.190 | 2.66 |
| 9 | 22.84 | 13116 | 15248.258 | 168 | 0.860 | 2.62 |
| 10 | 49.04 | 28666 | 22442.412 | 159 | 1.277 | 2.52 |
| 11 | 33.59 | 2116 | 244076.214 | 111 | 0.009 | 1.97 |
| 13 | 127.39 | 15332 | 3706.562 | 193 | 4.136 | 2.91 |
| 14 | 23.62 | 16623 | 106930.840 | 200 | 0.155 | 2.99 |
| 15 | 46.50 | 2070 | 96781.988 | 102 | 0.021 | 1.87 |
| 16 | 57.68 | 13535 | 15922.380 | 146 | 0.850 | 2.37 |
| 17 | 43.98 | 7496 | 533112.899 | 129 | 0.014 | 2.18 |
| 18 | 34.74 | 9162 | 9571.989 | 126 | 0.957 | 2.14 |

**Table 4.** Measured values captured using the iPhone (Size) and their similarity to the real measurements (Similarity) to evaluate the accuracy of iPhone cameras. The values were obtained by scanning rectangular boxes with sharp edges using the ‘Polycam App’ and comparing the dimensions to those measured manually with a meter.

| Object | Side | Size (cm) | Standard deviation (cm) | N | Real Size (cm) | Similarity (%) |
| --- | --- | --- | --- | --- | --- | --- |
| pink box | X | 27.4 | 0.34 | 3 | 25 | 90.4 |
| pink box | Y | 24.78 | 2.52 | 3 | 25 | 99.12 |
| pink box | Z | 27.4 | 0.49 | 3 | 25 | 90.4 |
| iPhone box | X | 9.68 | 0.06 | 3 | 9 | 92.44 |
| iPhone box | Y | 3.45 | 0.24 | 3 | 3 | 85 |
| iPhone box | Z | 17.32 | 0.27 | 3 | 16.5 | 95.03 |
| matches box | X | 7.05 | 0.2 | 3 | 6.5 | 91.54 |
| matches box | Y | 2.32 | 0.08 | 3 | 1.75 | 67.43 |
| matches box | Z | 11.44 | 0.12 | 3 | 11 | 96 |
| white box | X | 10.97 | 0.3 | 3 | 10 | 90.3 |
| white box | Y | 10.61 | 0.07 | 3 | 10 | 93.9 |
| white box | Z | 10.84 | 0.17 | 3 | 10 | 91.6 |

**Figure S1.**
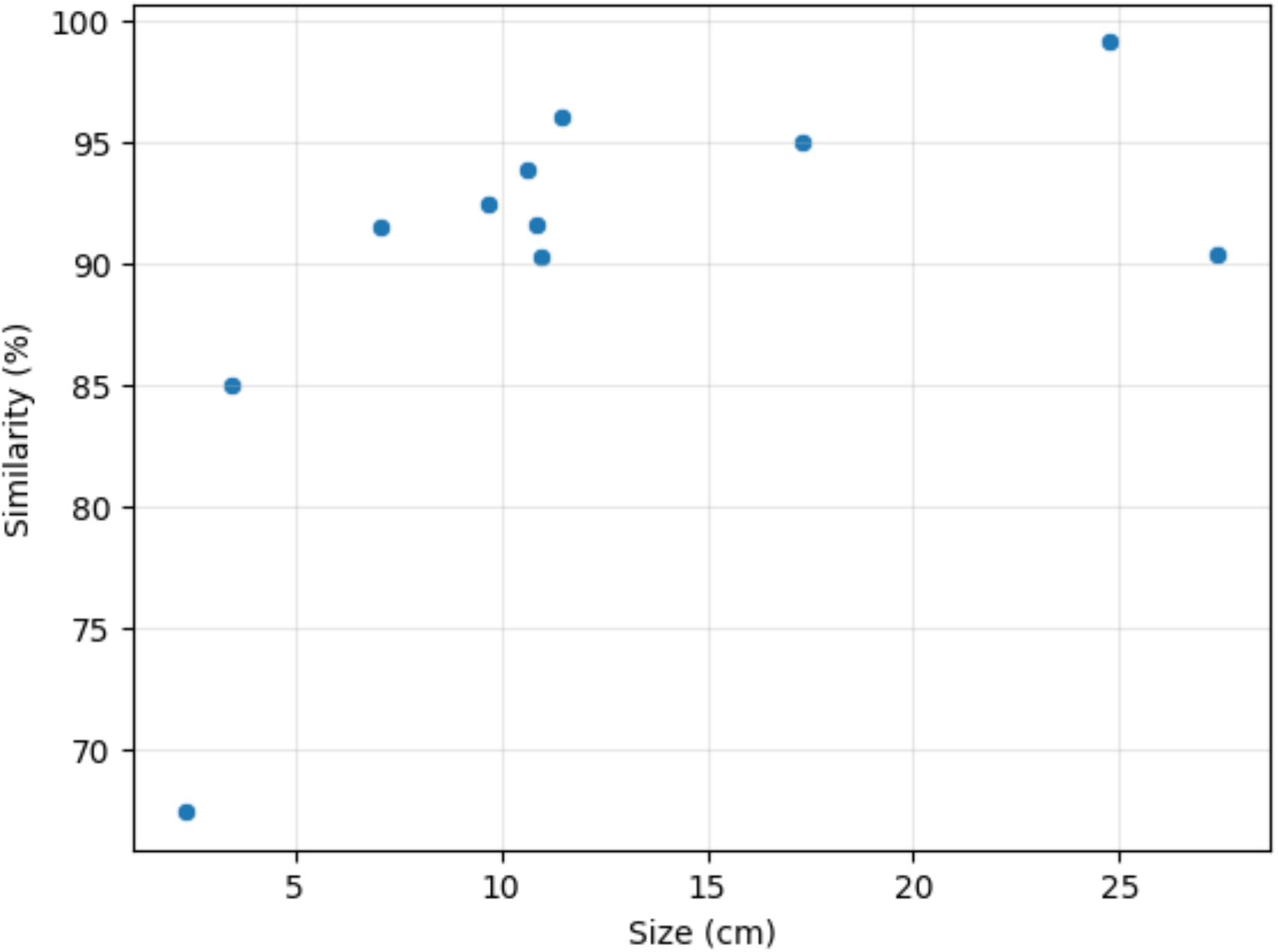
The percentage of similarity between the measured and real sizes for the hight, width and length dimensions of the rectangular boxes used for the iPhone’s technical test (Materials Table 4). Similarity increases as the size of the object dimension increases.

## Data Availability Statement

All statistical code and data files needed are available to download at https://github.com/gavrielbs/3D_Corn_Phenotype.

## Acknowledgments

This work was partially supported by a grant from the Hebrew University of Jerusalem Center for Interdisciplinary Data Science Research (CIDR). Menachem Moshelion received partial support through grants from the Horizon Europe Collaborative projects: EMPHASIS- GO (Grant Number 3012006413) and AGROSERV (Grant Number 3012006130). Matan Gavish was partially supported by the Israel Science Foundation (Grant No. 871/22).

**Disclaimer/Publisher’s Note:** The statements, opinions and data contained in all publications are solely those of the individual author(s) and contributor(s) and not of MDPI and/or the editor(s). MDPI and/or the editor(s) disclaim responsibility for any injury to people or property resulting from any ideas, methods, instructions or products referred to in the content.

## References

1. Kah, M.; Tufenkji, N.; White, J.C. Nano-Enabled Strategies to Enhance Crop Nutrition and Protection. Nat. Nanotechnol. 2019, 14, 532–540, doi:10.1038/s41565-019-0439-5.

2. Zhang, Q.; Ying, Y.; Ping, J. Recent Advances in Plant Nanoscience. Adv. Sci. 2022, 9, 2103414, doi:10.1002/advs.202103414.

3. Rosenzweig, C.; Iglesius, A.; Yang, X.B.; Epstein, P.R.; Chivian, E. Climate Change and Extreme Weather Events - Implications for Food Production, Plant Diseases, and Pests.

4. Piao, S.; Ciais, P.; Huang, Y.; Shen, Z.; Peng, S.; Li, J.; Zhou, L.; Liu, H.; Ma, Y.; Ding, Y.; et al. The Impacts of Climate Change on Water Resources and Agriculture in China. Nature 2010, 467, 43–51, doi:10.1038/nature09364.

5. Dall’erba, S.; Domínguez, F. The Impact of Climate Change on Agriculture in the Southwestern United States: The Ricardian Approach Revisited. Spat. Econ. Anal. 2016, 11, 46–66, doi:10.1080/17421772.2015.1076574.

6. Giraldo, J.P.; Wu, H.; Newkirk, G.M.; Kruss, S. Nanobiotechnology Approaches for Engineering Smart Plant Sensors. Nat. Nanotechnol. 2019, 14, 541–553, doi:10.1038/s41565-019-0470-6.

7. Zhang, C.; Kong, J.; Wu, D.; Guan, Z.; Ding, B.; Chen, F. Wearable Sensor: An Emerging Data Collection Tool for Plant Phenotyping. Plant Phenomics 5, 0051, doi:10.34133/plantphenomics.0051.

8. Improving Food Security through Increasing the Precision of Agricultural Development. In Precision Agriculture for Sustainability and Environmental Protection; Gassner, A., Coe, R., Sinclair, F., Eds.; Routledge, 2013; pp. 52–76 ISBN 978-0-203-12832-9.

9. Omia, E.; Bae, H.; Park, E.; Kim, M.S.; Baek, I.; Kabenge, I.; Cho, B.-K. Remote Sensing in Field Crop Monitoring: A Comprehensive Review of Sensor Systems, Data Analyses and Recent Advances. Remote Sens. 2023, 15, doi:10.3390/rs15020354.

10. Jin, S.; Sun, X.; Wu, F.; Su, Y.; Li, Y.; Song, S.; Xu, K.; Ma, Q.; Baret, F.; Jiang, D.; et al. Lidar Sheds New Light on Plant Phenomics for Plant Breeding and Management: Recent Advances and Future Prospects. ISPRS J. Photogramm. Remote Sens. 2021, 171, 202–223, doi:10.1016/j.isprsjprs.2020.11.006.

11. Su, Y.; Wu, F.; Ao, Z.; Jin, S.; Qin, F.; Liu, B.; Pang, S.; Liu, L.; Guo, Q. Evaluating Maize Phenotype Dynamics under Drought Stress Using Terrestrial Lidar. Plant Methods 2019, 15, 11, doi:10.1186/s13007-019-0396-x.

12. Zhou, L.; Gu, X.; Cheng, S.; Yang, G.; Shu, M.; Sun, Q. Analysis of Plant Height Changes of Lodged Maize Using UAV-LiDAR Data. Agriculture 2020, 10, doi:10.3390/agriculture10050146.

13. Underwood, J.P.; Hung, C.; Whelan, B.; Sukkarieh, S. Mapping Almond Orchard Canopy Volume, Flowers, Fruit and Yield Using Lidar and Vision Sensors. Comput. Electron. Agric. 2016, 130, 83–96, doi:10.1016/j.compag.2016.09.014.

14. Drusch, M.; Del Bello, U.; Carlier, S.; Colin, O.; Fernandez, V.; Gascon, F.; Hoersch, B.; Isola, C.; Laberinti, P.; Martimort, P.; et al. Sentinel-2: ESA’s Optical High-Resolution Mission for GMES Operational Services. Sentin. Missions - New Oppor. Sci. 2012, 120, 25–36, doi:10.1016/j.rse.2011.11.026.

15. Zhang, Z.; Boubin, J.; Stewart, C.; Khanal, S. Whole-Field Reinforcement Learning: A Fully Autonomous Aerial Scouting Method for Precision Agriculture. Sensors 2020, 20, doi:10.3390/s20226585.

16. Vadez, V.; Kholová, J.; Hummel, G.; Zhokhavets, U.; Gupta, S.K.; Hash, C.T. LeasyScan: A Novel Concept Combining 3D Imaging and Lysimetry for High-Throughput Phenotyping of Traits Controlling Plant Water Budget. J. Exp. Bot. 2015, 66, 5581–5593, doi:10.1093/jxb/erv251.

17. Luetzenburg, G.; Kroon, A.; Bjørk, A.A. Evaluation of the Apple iPhone 12 Pro LiDAR for an Application in Geosciences. Sci. Rep. 2021, 11, 22221, doi:10.1038/s41598-021-01763-9.

18. Zizka, A.; Joerger-Hickfang, T.; Imhof, S.; Méndez, L. LiDAR Sensors in Smartphones Can Enrich Herbarium Specimens with 3D Models of Habitat at High Precision and Little Cost. TAXON 2023, 72, 233–236, doi:10.1002/tax.12861.

19. Tatsumi, S.; Yamaguchi, K.; Furuya, N. ForestScanner: A Mobile Application for Measuring and Mapping Trees with LiDAR-Equipped iPhone and iPad. Methods Ecol. Evol. 2023, 14, 1603–1609, doi:10.1111/2041-210X.13900.

20. Vogt, M.; Rips, A.; Emmelmann, C. Comparison of iPad Pro®’s LiDAR and TrueDepth Capabilities with an Industrial 3D Scanning Solution. Technologies 2021, 9, doi:10.3390/technologies9020025.

21. Moshelion, M.; Dalal, A.; Shenhar, I.; Bourstein, R.; Mayo, A.; Grunwald, Y.; Averbuch, N.; Attia, Z.; Wallach, R. A Telemetric, Gravimetric Platform for Real-Time Physiological Phenotyping of Plant–Environment Interactions. J. Vis. Exp. 2020, e61280, doi:10.3791/61280.

22. Miao, T.; Wen, W.; Li, Y.; Wu, S.; Zhu, C.; Guo, X. Label3DMaize: Toolkit for 3D Point Cloud Data Annotation of Maize Shoots. GigaScience 2021, 10, giab031, doi:10.1093/gigascience/giab031.

23. Edelsbrunner, H.; Mücke, E.P. Three-Dimensional Alpha Shapes. ACM Trans Graph 1994, 13, 43–72, doi:10.1145/174462.156635.

24. Ao, Z.; Wu, F.; Hu, S.; Sun, Y.; Su, Y.; Guo, Q.; Xin, Q. Automatic Segmentation of Stem and Leaf Components and Individual Maize Plants in Field Terrestrial LiDAR Data Using Convolutional Neural Networks. Crop Phenotyping Stud. Appl. Crop Monit. 2022, 10, 1239–1250, doi:10.1016/j.cj.2021.10.010.

25. S. Jin; Y. Su; F. Wu; S. Pang; S. Gao; T. Hu; J. Liu; Q. Guo Stem–Leaf Segmentation and Phenotypic Trait Extraction of Individual Maize Using Terrestrial LiDAR Data. IEEE Trans. Geosci. Remote Sens. 2019, 57, 1336–1346, doi:10.1109/TGRS.2018.2866056.

26. Forero, M.G.; Murcia, H.F.; Méndez, D.; Betancourt-Lozano, J. LiDAR Platform for Acquisition of 3D Plant Phenotyping Database. Plants 2022, 11, doi:10.3390/plants11172199.

27. Paulus, S. Measuring Crops in 3D: Using Geometry for Plant Phenotyping. Plant Methods 2019, 15, 103, doi:10.1186/s13007-019-0490-0.

28. Alam, M.S.; Lamb, D.W.; Warwick, N.W.M. A Canopy Transpiration Model Based on Scaling Up Stomatal Conductance and Radiation Interception as Affected by Leaf Area Index. Water 2021, 13, doi:10.3390/w13030252.

